# Loss of the MeCP2 gene in parvalbumin interneurons leads to an inhibitory deficit in the amygdala and affects its functional connectivity

**DOI:** 10.1101/2024.05.30.596683

**Authors:** Maj Liiwand, Joni Haikonen, Bojana Kokinovic, Svetlana M. Molchanova, Teemu Aitta-aho, Sari E. Lauri, Maria Ryazantseva

## Abstract

**Background:** *MECP2* gene is located in the X-chromosome and encodes a methyl-CpG-binding protein involved in transcription regulation. The loss-of-function mutation of the *MECP2* gene, leads to severe neurodevelopmental syndrome, Rett syndrome. Clinical picture of Rett syndrome includes, among other symptoms, social deficits and heightened anxiety. The amygdala is involved in the regulation of social behavior as well as fear and anxiety. Here, we investigated the effect of the *MeCP2* gene ablation in the parvalbumin interneurons on the microcircuit and functional connectivity of the amygdala.

**Methods:** Males with conditional knock-out of the *MeCP2* gene in the parvalbumin interneurons were used as a genetic mouse model of MeCP2 loss in parvalbumin interneurons. Littermates with the intact gene were used as controls. Ex-vivo brain slice electrophysiology, combined with pharmacology and optogenetics, was used to characterize microcircuits within the lateral amygdala. Synaptic currents and excitability of parvalbumin interneurons and principal neurons were analyzed by a whole-cell patch clamp. In-vivo functional ultrasound was used to visualize the connectivity within the amygdala–ventral hippocampus–prefrontal cortex triad.

**Results:** Loss of *MeCP2* in parvalbumin interneurons significantly reduced the GABAergic synaptic input to the principal neurons in the lateral amygdala. The decreased inhibitory drive was accompanied by an increase in the excitability of principal neurons in the lateral amygdala. The in vivo functional connectivity of the amygdala-ventral hippocampus and amygdala-prefrontal cortex was significantly reduced in conditional knock-outs compared to their littermates with the intact gene in the X chromosome.

**Conclusions:** Our study characterized the consequences of *MeCP2* gene loss in the parvalbumin interneurons on the amygdala connectivity and microcircuit and provided evidence supporting the previous findings on the role of interneurons in the functional deficit observed in animal models with *MeCP2* loss.

## Background

The *Methyl CpG binding protein 2 (MECP2)* gene, located on the X-chromosome is associated with neurodevelopmental syndromes: Rett syndrome and MECP2 duplication syndrome [1, 2]. Both syndromes significantly affect brain development, with symptoms emerging early in life. In addition, there is evidence connecting mutations in the *MECP2* gene to autism spectrum disorders [3]. Overall, patients carrying genetic alterations in *MECP2* are anxious and have poor social interaction skills and learning impairment [1–4]. The biology behind these symptoms is poorly understood.

Animal studies demonstrated that the majority of features, including social and learning deficits, can be replicated in mice by depleting the *MeCP2* gene in inhibitory neurons [5, 6]. Moreover, further studies showed that the conditional knockout of the gene in parvalbumin (PV) type of interneurons leads to motor, sensory, social, and cognitive alterations [7]. The conditional knock-out of *MeCP2* in PV interneurons replicates deficiency in the cortical plasticity and related behavioral features of mice, which lack one functional allele of *MeCP2* [8]. Although these results stress the importance of MeCP2 protein for the function of inhibitory neurons and connect the neurological consequences to the compromised GABAergic signaling, they do not provide a full picture of the circuit deficiency in the affected brain areas.

The amygdala is heavily implicated in social interaction [9–12]. It is suggested that the complex network of reciprocally interconnected structures, including amygdala, the medial prefrontal cortex and ventral hippocampus (mPFC and vHPC), is crucial for regulating social interaction [10, 12, 13]. Basolateral (BLA) amygdala-projecting neurons in the infralimbic cortex (IL) are preferentially activated in response to a social cue, as compared to BLA-projecting neurons in the prelimbic cortex (PL). Chemogenetic examination of these sub-circuits in mice shows that activation of PL-BLA or inhibition of IL-BLA circuits impairs social behavior [12]. BLA inputs to the vHPC are capable of modulating social behaviors in a bidirectional manner: inhibition of BLA-vHPC projections significantly increases social interaction, while activation of the projections leads to the opposite effect [10]. We hypothesized that the functional connectivity of the amygdala with brain areas involved in regulating social interaction is affected in mice lacking MeCP2 in PV interneurons.

Mice lacking the *MeCP2* gene in the PV interneurons demonstrated a learning and memory deficit in fear conditioning [7]. Fear learning is supported by BLA excitatory projection neurons, and their activity is regulated by potent perisomatic inhibition provided by PV interneurons. Silencing PV interneurons during the US augments fear learning, whereas activating them reduces fear learning [14, 15]. Only PV interneurons in the lateral (LA) part of BLA, but not the basal part (BA), possess complex dendritic arborizations, receive potent excitatory drive, and mediate feedforward inhibition onto principal neurons [16]. After fear conditioning, they exhibit nucleus- and target-selective plasticity, resulting in a persistent reduction of their excitatory input and inhibitory output in LA but not BA [16]. Previously, the knock-out of the *MeCP2* gene in other brain areas has been demonstrated to affect the excitatory input to [17, 18] and excitability [19] of PV interneurons. We hypothesized that the loss of MeCP2 influences excitatory or inhibitory synaptic connectivity of PV interneurons and leads to improper inhibition provided to principal neurons in LA.

Here, we compare the synaptic and excitatory properties of PV interneurons of LA lacking the *MeCP2* gene to those of PV interneurons of littermate mice with the intact gene. In addition, we investigate the properties of excitatory principal neurons in LA of these mice and test for fast and slow deficits in GABAergic inhibition. Moreover, we test the functional connectivity of the amygdala in vivo to detect the consequences of genetic ablation of the *MeCP2* gene in PV interneurons for BLA-vHPC and BLA-mPFC circuits. Overall, we provide new evidence of the role of the *MeCP2* gene in PV interneurons function in regulating amygdala local inhibition and functional connectivity.

## Results

### PV neurons lacking the MeCP2 gene receive elevated excitatory input but release less GABA to principal neurons in the lateral amygdala

We used a previously published strategy for Cre-dependent conditional knock-out of the floxed MeCP2 gene in the PV interneurons of mouse brains [19]. PV-Cre males were crossed with MeCP2^fl/wt^ females to obtain littermate males with conditional knock-out (cKO) of the MeCP2 gene in PV interneurons (PV-Cre::MeCP2^fl/y^) and males with an intact MeCP2 gene (PV-Cre::MeCP2^wt/y^, which will be referred to as PV-Cre for simplicity). PV-Cre littermates were used as age-matched control for PV-Cre::MeCP2^fl/y^ in all experiments, and mice were used as adults (1.5-2 months old). To visualize the PV interneurons in the LA nucleus of the amygdala, mice were locally injected with AAV construct, encoding Cre-dependent EGFP. Synaptic properties and excitability of PV interneurons were then addressed in acute coronal slices containing BLA by whole-cell patch clamp recording. Spontaneous excitatory postsynaptic currents (sEPSC) from EGFP-labelled neurons in LA were recorded as inward events when the recording was done at −50 mV holding potential and with a low chloride pipette solution (Figure 1 a.). The frequencies of sEPSC were not different for PV interneurons among genotypes, while the amplitudes of sEPSC were significantly increased in PV interneurons lacking the MeCP2 gene (freq.: 29.41 ± 4.823 Hz for PV-Cre vs. 35.87 ± 6.735 Hz for PV-Cre::MeCP2^fl/y^, t-test, t=0.7666, df=21, p=0.4519; ampl.: 20.90 ± 1.510 pA for PV-Cre vs. 28.83 ± 3.015 pA, t-test, t=2.286, df=21, *p=0.0327) (Figure 1 a.). The spontaneous inhibitory postsynaptic currents (sIPSC) were recorded simultaneously as outward currents at −50 mV holding potential. No differences were observed for the frequencies or amplitudes of sIPSC in PV interneurons among the genotypes (freq.: 13.25 ± 2.810 Hz for PV-Cre vs. 17.66 ± 2.966 Hz for PV-Cre::MeCP2^fl/y^, t-test, t=1.075, df=21, p=0.2948; ampl.: 13.07 ± 0.6627 for PV-Cre vs. 14.83 ± 0.9525 for PV-Cre::MeCP2^fl/y^, t-test, t=1.542, df=21, p=0.1381). To further address the GABA release from the PV interneurons to principal neurons in LA, the AAV viral vectors encoding Cre-dependent channelrhodopsin-2 was locally injected into BLA. Blue light was used to induce synaptic release of GABA from PV interneurons. IPSC were recorded in whole-cell mode from principal neurons in LA with a high chloride pipette solution and −80 mV holding potential. NBQX and AP-5 in ASCF were used to block AMPA and NMDA currents correspondingly. A paired-pulse protocol was applied to address the probability of GABA release. The paired-pulse ratio of IPSC in principal neurons in LA of PV-Cre mice was significantly lower than in principal neurons in LA of PV-Cre::MeCP2^fl/y^ mice as compared to controls (PPR: 0.8074 ± 0.04814 for PV-Cre vs. 1.117 ± 0.1040 for PV-Cre::MeCP2^fl/y^, t-test, t=2.349, df=20, *p=0.0292) (Figure 1. b.), indicating lower GABA release from PV interneurons lacking the MeCP2 gene.

**Figure 1.**
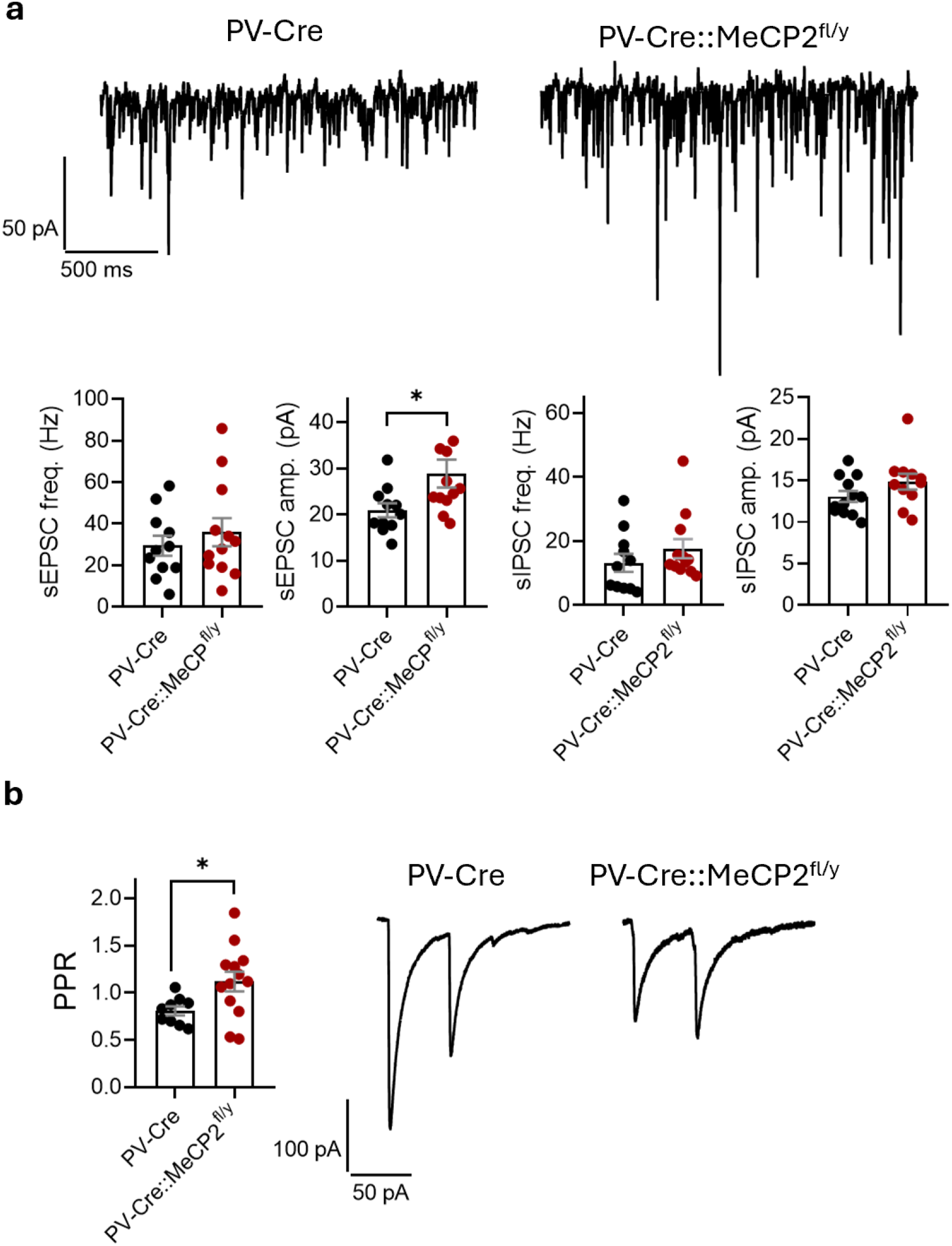
Lack of MeCP2 gene in the parvalbumin interneurons (PV) of the lateral amygdala leads to post-synaptic and pre-synaptic changes. **a**. Example traces of whole-cell recordings of synaptic currents at –50 mV holding potential done in the PV+ interneurons within lateral amygdala of PV-Cre and PV-Cre::MeCP2^fl/y^ littermates (n=11 neurons from 3 mice for PV-Cre; n=12 neurons from 3 mice for PV-Cre::MeCP2^fl/y^). Frequencies and amplitudes of spontaneous excitatory postsynaptic currents (sEPSC) (freq.: t-test, t=0.7666, df=21, p=0.4519; ampl.: t-test, t=2.286, df=21, *p=0.0327). Frequencies and amplitudes of spontaneous inhibitory postsynaptic currents (sIPSC) (freq.: t-test, t=1.075, df=21, p=0.2948; ampl.: t-test, t=1.542, df=21, p=0.1381). **b**. Paired pulse ratios of inhibitory synaptic responses recorded from principal neurons within the lateral amygdala of PV-Cre and PV-Cre::MeCP2^fl/y^ littermates Cre-dependently expressing channeloropsin-2 in the PV interneurons (n=9 neurons from 4 mice for PV-Cre; n=13 neurons from 3 mice for PV-Cre::MeCP2^fl/y^; t-test, t=2.349, df=20, *p= 0.0292). All the data is shown as mean ± S.E.M.

### Loss of the MeCP2 gene in PV interneurons within LA does not affect their excitability

The EGFP-labelled PV interneurons in LA were examined for their passive membrane properties and intrinsic excitability by intracellular injection of step currents in the current-clamp mode. There was no difference between the genotypes for action potential firing frequencies (two-way ANOVA, genotype effect: F(1,15)=0.1802, p=0.6773) (Figure 2. a), resting membrane potential (−58.11 ± 2.732 mV for PV-Cre vs. −52.89 ± 3.821 mV for PV-Cre::MeCP2^fl/y^, t-test, t=1.086, df=15, p=0.2945), rheobase (102.2 ± 15.90 pA for PV-Cre vs. 88.33 ± 9.789 pA for PV-Cre::MeCP2^fl/y^, t-test, t=0.7438, df=16, p=0.4678), action potential half-width (0.3850 ± 0.01626 ms for PV-Cre vs. 0.3944 ± 0.02249 ms for PV-Cre::MeCP2^fl/y^, t-test, t=0.3326, df=15, p=0.7440), and firing threshold (41.00 ± 1.345 mV for PV-Cre vs. 40.80 ± 1.929 mV for PV-Cre::MeCP2^fl/y^, t-test, t=0.08054, df=15, p= 0.9369) (Figure 2 b.).

**Figure 2.**
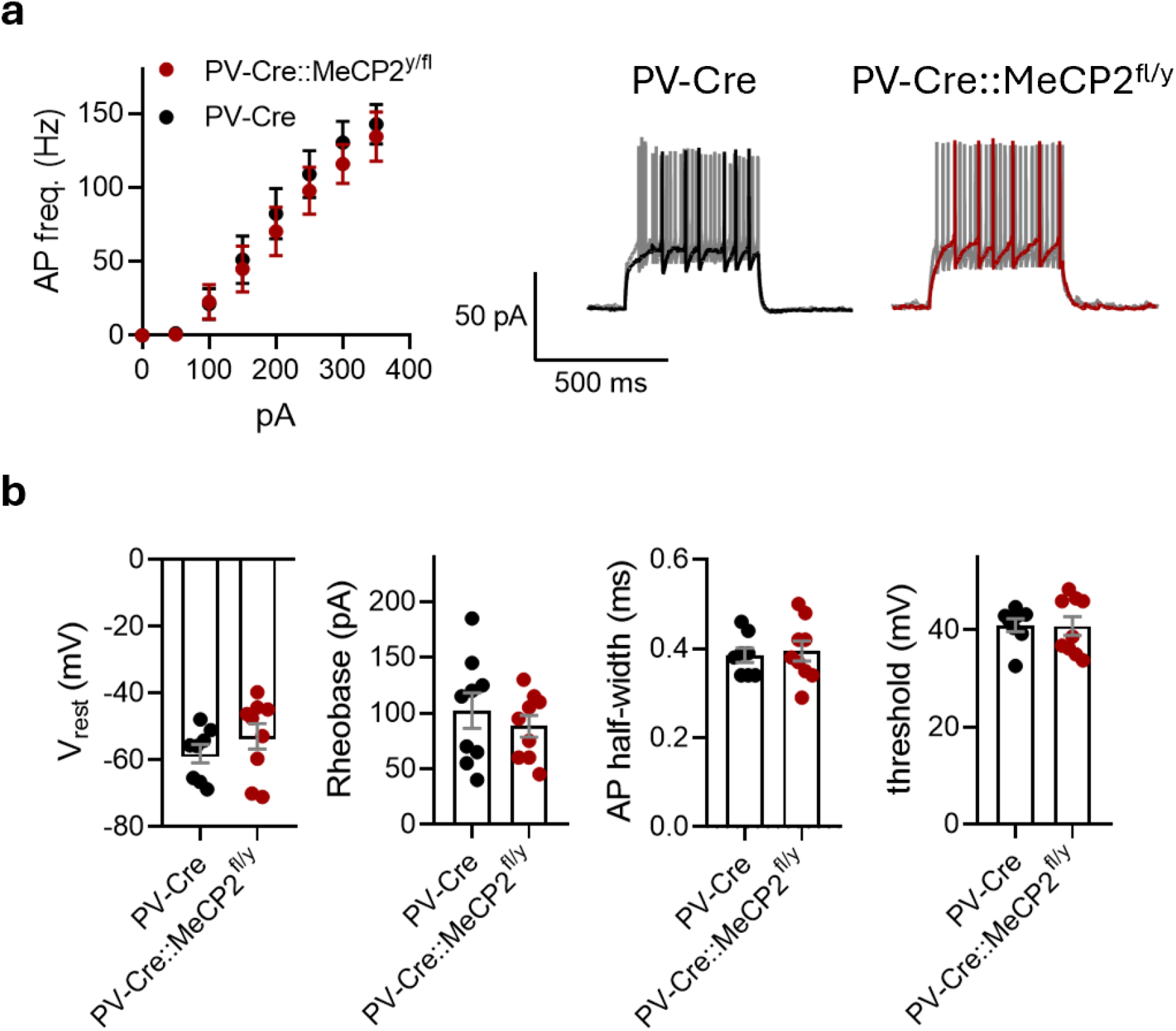
Lack of MeCP2 gene in the parvalbumin interneurons (PV) of the lateral amygdala does not affect their excitability. **a**. Action potential (AP) firing frequencies in response to current steps plotted from the rheobase with 25 pA increment (n=8 neurons from 3 mice for PV-Cre; n=9 neurons from 3 mice for PV-Cre::MeCP2^fl/y^; two-way ANOVA, genotype effect: F(1, 15) = 0.1802, p= 0.6773) and example traces of AP burst in response to 25 and 100 pA current steps. **b**. Resting membrane potential (V_rest_), rheobase, AP half-width and firing threshold for PV interneurons in the lateral amygdala of PV-Cre and PV-Cre::MeCP2^fl/y^ littermates (n=8 neurons from 3 mice for PV-Cre; n=9 neurons from 3 mice for PV-Cre::MeCP2^fl/y^; V_rest_: t-test, t=1.086, df=15, p= 0.2945; rheobase: t-test, t=0.7438, df=16, p=0.4678; AP half-width: t-test, t=0.3326, df=15, p= 0.7440; threshold: t-test, t=0.08054, df=15, p= 0.9369). All the data is shown as mean ± S.E.M.

### Principal neurons in LA of mice lacking MeCP2 in the PV interneurons demonstrate increased excitability

Principal neurons in LA are biological substrates for plasticity and associative learning. Their excitability plays a significant role in plasticity and memory trace allocation and is subject to changes in fear conditioning [20]. Principal neurons in LA of PV-Cre::MeCP2^fl/y^ mice had an increase in the action potential firing frequencies recorded with the application of step currents in the current-clamp mode (two-way ANOVA, genotype effect: F(1, 29) = 5.217, *p=0.0299) (Figure 3. a.). At the same time, there were no differences in the resting membrane potential (−59.82 ± 1.096 mV for PV-Cre vs. −60.00 ± 1.701 mV for PV-Cre::MeCP2^fl/y^, t-test, t=0.09421, df=30, p=0.9256), amplitude of firing threshold (39.87 ± 1.432 mV for PV-Cre vs. 37.29 ± 1.227 mV for PV-Cre::MeCP2^fl/y^, t-test, t=1.300, df=29, p=0.2038), action potential half-width (1.609 ± 0.1302 ms for PV-Cre vs. 1.389 ± 0.06622 ms PV-Cre::MeCP2^fl/y^, t-test, t=1.373, df=28, p=0.1806) or rheobase (66.67 ± 8.556 pA for PV-Cre vs. 66.92 ± 6.241 pA for PV-Cre::MeCP2^fl/y^, t-test, t=0.02248, df=29, p=0.9822) (Figure 3. b.).

**Figure 3.**
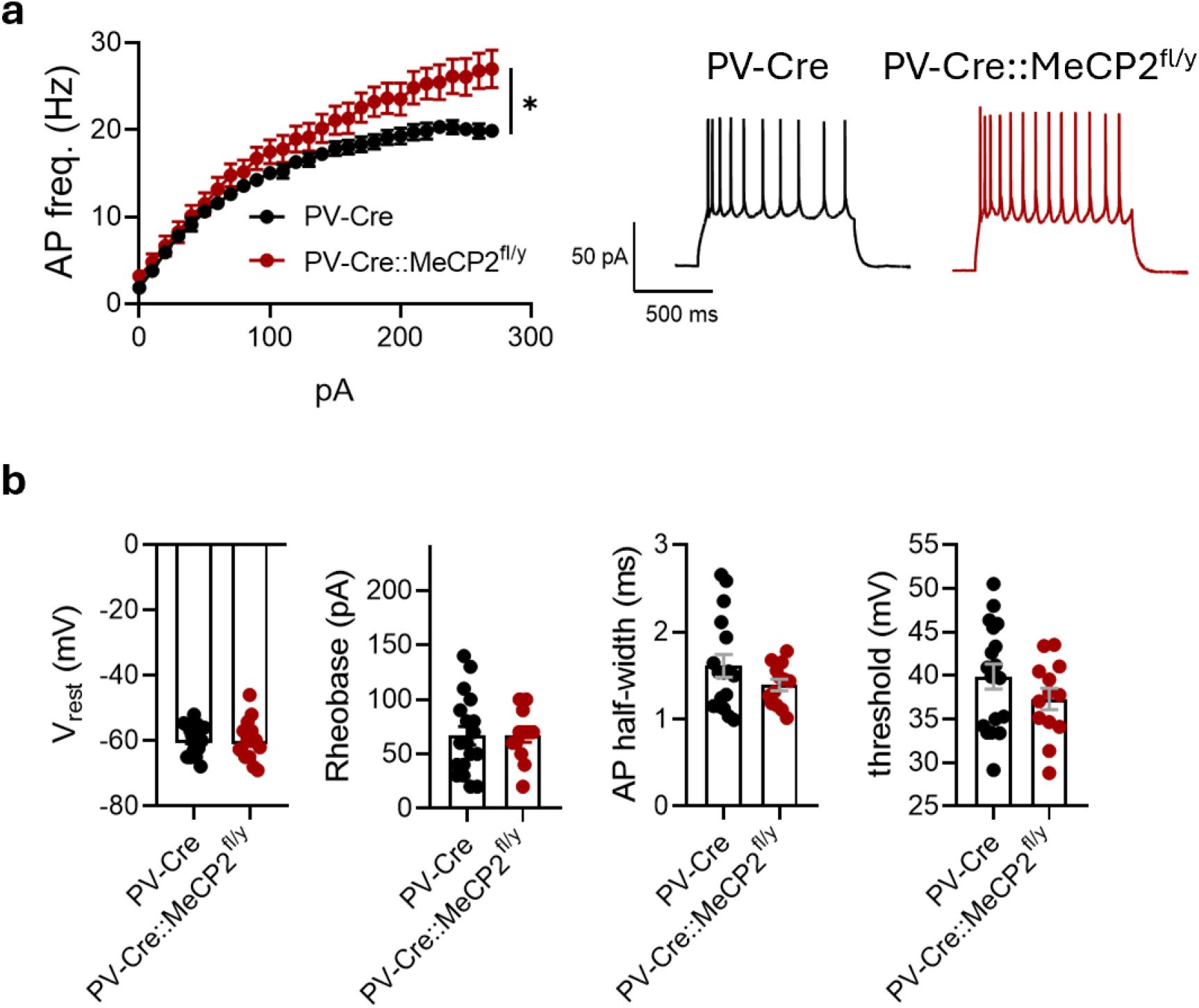
Lack of the MeCP2 gene in PV interneurons increases the excitability of principal neurons in the lateral amygdala. **a**. Action potential (AP) firing frequencies of principal neurons in response to current steps plotted from the rheobase with 10 pA increment (n=18 neurons from 5 mice for PV-Cre; n=13 neurons from 3 mice for PV-Cre::MeCP2^fl/y^; two-way ANOVA, genotype effect: F(1, 29) = 5.217, *p=0.0299) and example traces of AP burst in response to 150 pA current step. **b**. Resting membrane potential (V_rest_), rheobase, action potential (AP) half-width and firing threshold for principal neurons in the lateral amygdala of PV-Cre and PV-Cre::MeCP2^fl/y^ littermates (n=18 neurons from 5 mice for PV-Cre; n=13 neurons from 3 mice for PV-Cre::MeCP2^fl/y^; V_rest_: t-test, t=0.09421, df=30, p=0.9256; rheobase: t-test, t=0.02248, df=29, p=0.9822; AP half-width: t-test, t=1.373, df=28, p= 0.1806; threshold: t-test, t=1.300, df=29, p=0.2038). All the data is shown as mean ± S.E.M.

### Loss of MeCP2 in PV interneurons leads to an inhibitory deficit in LA, affecting both fast and slow inhibition

PV interneurons potently modulate the excitability of principal neurons in the amygdala through perisomatic innervation. sIPSC and sEPSC were recorded simultaneously in whole-cell mode with a low chloride pipette solution at −50 mV holding potential from principal neurons of LA. There were no differences in the amplitudes or frequencies of sEPSC between genotypes (freq.: 7.156 ± 1.893 Hz for PV-Cre vs. 3.662 ± 0.8540 Hz for PV-Cre::MeCP2^fl/y^, t-test, t=1.783, df=20, p= 0.0897; ampl.: 15.02 ± 0.9445 pA for PV-Cre vs. 15.00 ± 1.082 for PV-Cre::MeCP2^fl/y^, t-test, t=0.01377, df=20, p=0.9891) (Figure 4. a.). At the same time, a significantly lower frequency of sIPSC was detected for principal neurons in LA of cKO mice which aligns with the reduction of GABA release from PV interneurons (freq.: 4.399 ± 0.7893 Hz for PV-Cre vs. 1.492 ± 0.4179 Hz for PV-Cre::MeCP2^fl/y^, t-test, t=3.470, df=21, **p= 0.0023) (Figure 4. a.). The amplitudes of sIPSC were the same (amp.: 16.12 ± 0.9511 pA for PV-Cre vs. 14.83 ± 0.8562 pA for PV-Cre::MeCP2^fl/y^, t=1.009, df=20, p=0.3250) (Figure 4. a.). The lack of changes in sEPSC due to MeCP2 cKO was additionally supported by the absence of visible alterations in the densities of the dendritic spines at principal neurons filled with biocytin during recording and analyzed post-hoc (0.9852 ± 0.08029 spine/μM for PV-Cre vs. 0.8314 ± 0.08733 spine/μM for PV-Cre::MeCP2^fl/y^, Mann-Whitney test, U=70, p=0.3255) (Figure 4. b.). This indicates that the loss of MeCP2 in PV interneurons does not affect the excitatory postsynaptic connectivity of principal neurons in LA while mainly reducing their inhibitory input.

**Figure 4.**
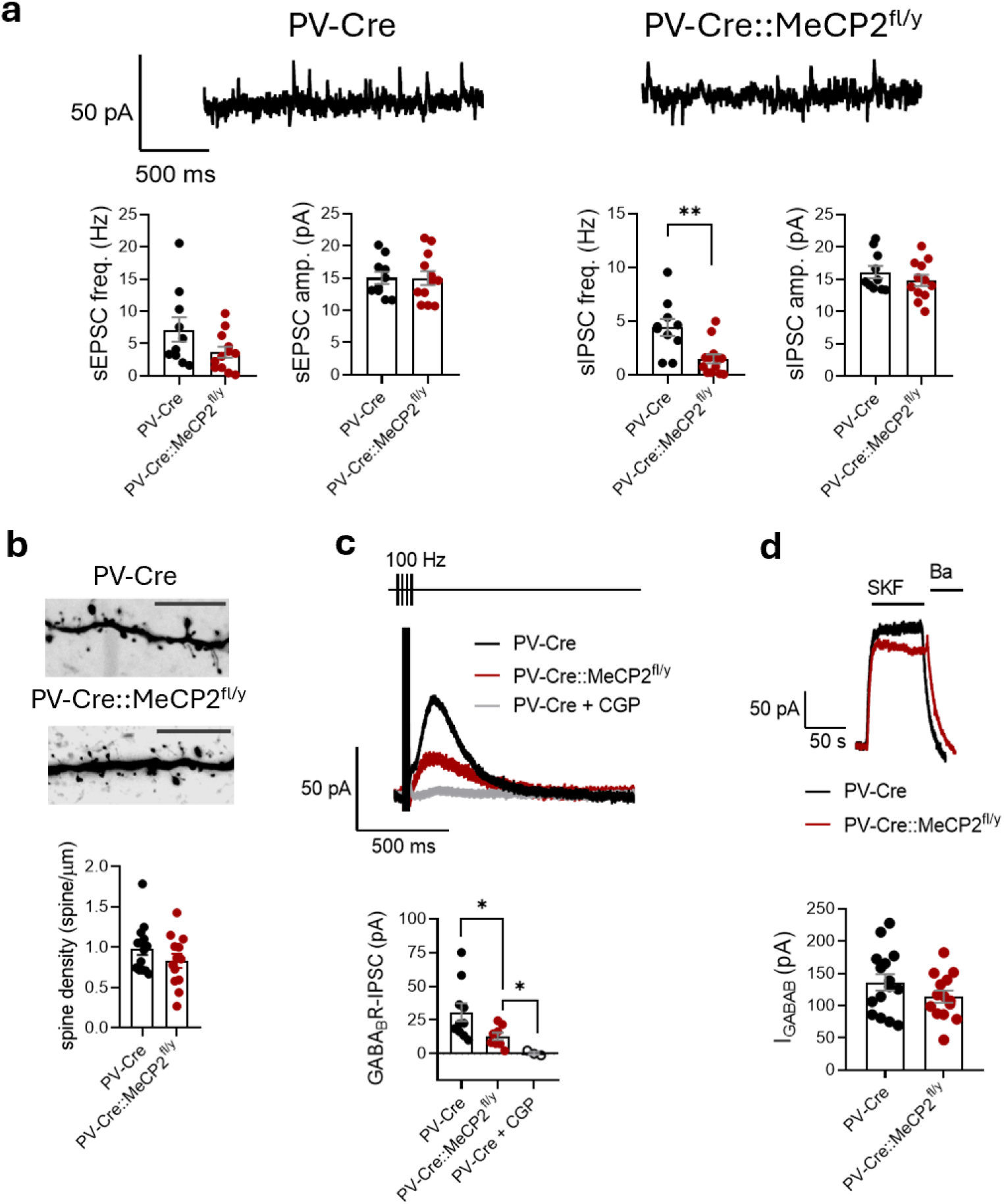
Lack of the MeCP2 gene in PV interneurons reduces fast and slow inhibitory input to the principal neurons in the lateral amygdala. **a**. Example traces of whole-cell recordings of synaptic currents at –50 mV holding potential done in the principal neurons within the lateral amygdala of PV-Cre and PV-Cre::MeCP2^fl/y^ littermates (n=10 neurons from 4 mice for PV-Cre; n=12 neurons from 3 mice for PV-Cre::MeCP2^fl/y^). Frequencies and amplitudes of sEPSC (freq.: t-test, t=1.783, df=20, p= 0.0897; ampl.: t-test, t=0.01377, df=20, p=0.9891). Frequencies and amplitudes of sIPSC (freq.: t-test, t=3.470, df=21, **p= 0.0023; ampl.: t-test, t=1.009, df=20, p=0.3250). **b**. Spine densities of principal neurons in the lateral amygdala (n=14 neurons from 4 mice for PV-Cre; n=13 neurons from 3 mice for PV-Cre::MeCP2^fl/y^; Mann-Whitney test, U=70, p=0.3255). Example images of spines for both genotypes, scale bar is 10 μm. **c**. Amplitudes of stimulation-induced slow inhibitory postsynaptic currents in principal neurons of lateral amygdala recorded at −50 mV holding potential in the presence of NBQX (50 μM), AP-5 (50 μM), and PiTx (100 μM). 5 μM CGP 55845 (CGP) was applied to block GABA_B_R responses of the principal neurons of PV-Cre mice. (n=10 neurons from 3 mice for PV-Cre; n=8 neurons from 3 animals for PV-Cre::MeCP2^fl/y^; n=3 neurons from 3 mice for CGP application; PV-Cre vs. PV-Cre::MeCP2^fl/y^: t-test, t=2.277, df=16, *p=0.0368; PV-Cre::MeCP2^fl/y^ vs. PV-Cre + CGP: t-test, t=2.662, df=9, *p=0.0260). Example traces of slow IPSCs in response to high-frequency stimulation by 4 pulses with 10 ms interevent interval. **d**. Amplitudes of outward currents induced by application of selective GABA_B_R agonist SKF 97541 (25 μM) and recorded at −50 mV holding potential. Recordings were done in the presence of NBQX (50 μM), AP-5 (50 μM), and PiTx (100 μM). (n=15 neurons from 3 mice for PV-Cre; n=14 neurons from 3 animals for PV-Cre::MeCP2^fl/y^; t-test, t=1.372, df=27, p=0.1814). Example traces of currents induced by SKF and sensitive to 2 mM of barium chloride (Ba). All the data is shown as mean ± S.E.M.

In addition to fast synaptic inhibition through the GABA_A_ receptors, the principal neurons in LA receive slow inhibition by the postsynaptic metabotropic GABA_B_-receptors that activate G protein-coupled inwardly rectifying potassium channels (GIRKs) [21]. To assess the slow inhibition, the release of GABA in the brain slices was activated with the stimulating electrode placed at the upper corner of the cortical side of LA. The recording was done from principal neurons in the middle of LA in the presence of NBXQ, AP-5, and picrotoxin to block AMPA, NMDA, and GABA_A_ currents, respectively, and the pipette contained a high potassium solution. Repetitive electrical stimuli (4 stimuli at 100 Hz) elicited a robust GABA_B_ current at - 50 mV holding potential. The current was sensitive to a potent inhibitor of GABA_B_-receptor CGP 55845 (Figure 4 c.). The maximum amplitude of the evoked GABA_B_ current was lower in principal neurons of Cre::MeCP2^fl/y^ mice than in controls (30.69 ± 6.649 pA for PV-Cre vs. 12.77 ± 2.743 pA for PV-Cre::MeCP2^fl/y^, t-test, t=2.277, df=16, *p=0.0368) (Figure 4. c.). Fast application of the potent GABA_B_-receptor agonist SKF 97541 [22], however, induced comparable amplitudes of agonist-induced currents in both genotypes (135.9 ± 12.76 pA for PV-Cre vs. 113.9 ± 9.384 pA for PV-Cre::MeCP2^fl/y^, t-test, t=1.372, df=27, p=0.1814). The data indicate that the reduction in GABA_B_ currents aligns with the deficit of the inhibition provided by interneurons to principal neurons in LA of mice lacking MeCP2 in PV interneurons. As the potent GABA_B_-receptor agonist can still induce the current of amplitude comparable to controls, there should be enough receptors and GIRK channels to provide slow inhibition when the GABA signal is sufficient.

### Loss of MeCP2 in PV interneurons affects functional connectivity in BLA-vHPC and BLA-mPFC circuits in vivo at resting state

Functional ultrasound (fUS) imaging has broad spatial coverage, providing large-scale measurements of brain activity in vivo [23]. Functional connectivity of the LA, BA and basomedial (BMA) amygdala nuclei with ventral hippocampus (vHPC), PL and IL was analysed simultaneously by fUS imaging in anesthetized mice. The connectivity was assayed in coronal planes, based on the calculation of mean Pearson correlation factor in the fUS signal between the anatomically defined regions of interest [24]. The correlation coefficients were then Fisher-transformed and used to build the functional connectivity matrices and statistical analysis. The results demonstrated a general trend for reduction of the amygdala connectivity with regions of interest in the brain of cKO mice (Figure 5. a.). Compared to control littermates, the correlation in activity was significantly reduced between BA nucleus and vHPC (0.6272 ± 0.1720 for PV-Cre vs. 0.2570 ± 0.06707 for PV-Cre::MeCP2^fl/y^, multiple t-tests, Holm-Sidak, t ratio=2.941, df=36.00, *p=0.016960) (Figure 5. b.) as well as with both PL and IL (PV-Cre vs. PV-Cre::MeCP2^fl/y^: for PL-BA, 0.1477 ± 0.05233 vs. 0.01462 ± 0.02987, multiple t-tests, Holm-Sidak, t=3.089, df=48.00, *p=0.01327; for IL-BA, 0.1650 ± 0.05335 vs. 0.04040 ± 0.01749, multiple t-tests, Holm-Sidak, t=2.967, df=48.00, *p=0.01395) (Figure 5. c. and d.). Interestingly, when the connectivity between vHPC and mPFC was evaluated, the reduction was significant only in vHPC-IL but not in vHPC-PL regions (PV-Cre vs. PV-Cre::MeCP2^fl/y^: for PL-vHPC, 0.1634 ± 0.05447 vs. 0.09774 ± 0.02235, multiple t-tests, Holm-Sidak, t=1.524, df=48.00, p=0.3505; for IL-vHPC, 0.2345 ± 0.04655 vs. 0.06915 ± 0.02221, multiple t-tests, Holm-Sidak, t=3.940, df=48.00, **p=0.0010) (Figure 5. c. and d.).

**Figure 5.**
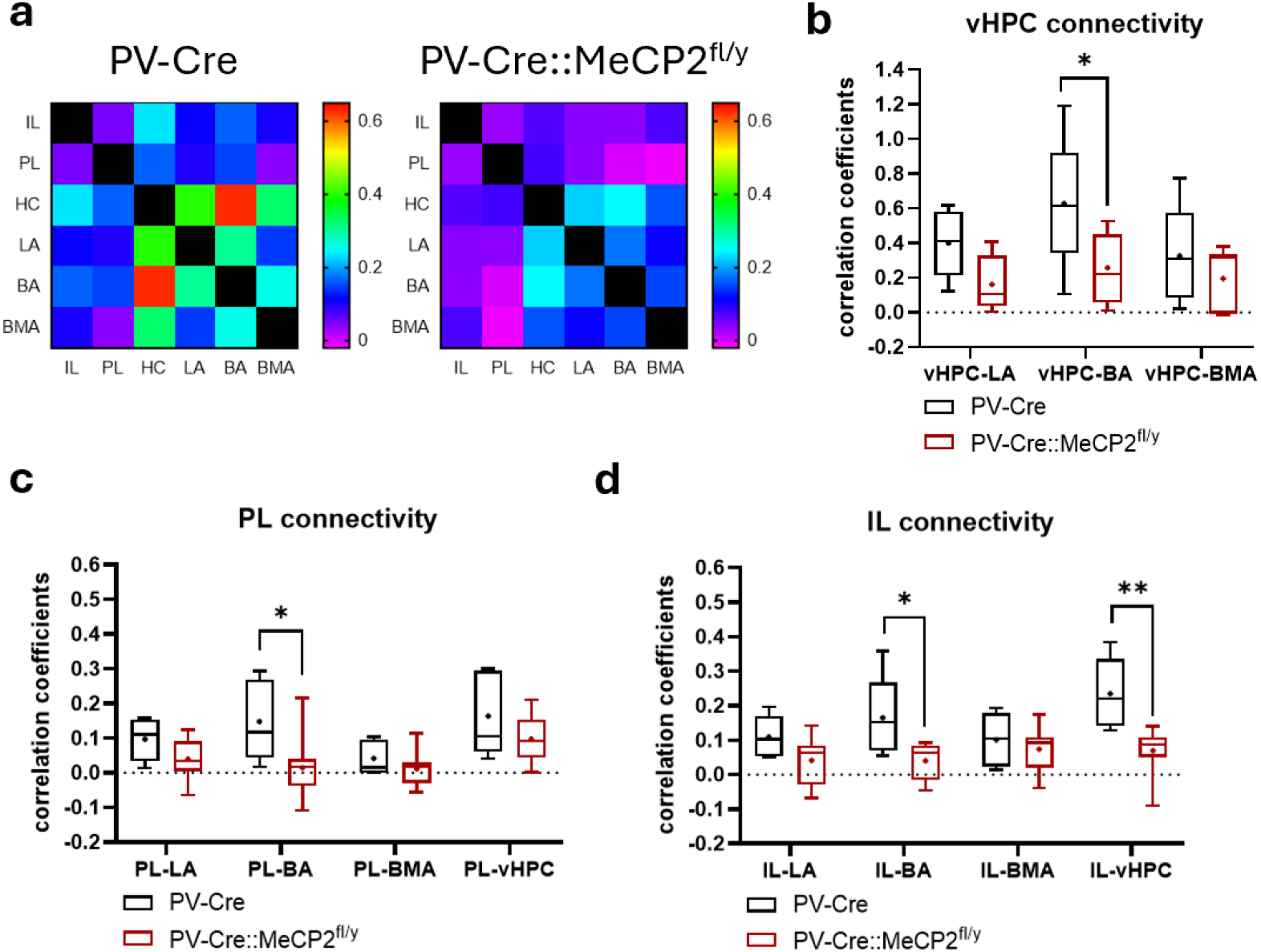
Lack of the MeCP2 gene in the PV interneurons affects functional connectivity between the amygdala, ventral hippocampus, and prefrontal cortex. **a**. Heat-maps of Pearson correlation coefficients of activity between following brain areas of anesthetized mice with PV-Cre and PV-Cre::MeCP2^fl/y^ genotypes: infralimbic (IL) and prelimbic (PL) medial prefrontal cortex, ventral hippocampus (vHPC), lateral amygdala (LA), basal amygdala (BA), basal medial amygdala (BMA). **b**. Comparison of the Pearson correlation coefficients between activity in vHPC and amygdala of PV-Cre and PV-Cre::MeCP2^fl/y^ mice (n=5 mice for PV-Cre, n=9 mice for PV-Cre::MeCP2^fl/y^; multiple t-tests, Holm-Sidak: vHPC-LA: t=1.894, df=36.00, p=0.1282; vHPC-BA: t=2.941, df=36.00, *p=0.0169; vHPC-BMA: t=1.039, df=36.00, p=0.3059). **c**. Comparison of the Pearson correlation coefficients between activity in PL and amygdala of PV-Cre and PV-Cre::MeCP2^fl/y^ mice (n=5 mice for PV-Cre, n=9 mice for PV-Cre::MeCP2^fl/y^; multiple t-tests, Holm-Sidak: PL-LA: t=1.338, df=48.00, p=0.3505; PL-BA: t=3.089, df=48.00, *p=0.01327; PL-BMA: t=0.7150, df=48.00, p=0.4780, PL-vHPC: t=1.524, df=48.00, p=0.3505). **d**. Comparison of the Pearson correlation coefficients between activity in IL and amygdala of PV-Cre and PV-Cre::MeCP2^fl/y^ mice (n=5 mice for PV-Cre, n=9 mice for PV-Cre::MeCP2^fl/y^; multiple t-tests, Holm-Sidak: IL-LA: t=1.619, df=48.00, p=0.2114; IL-BA: t=2.967, df=48.00, *p=0.01395; IL-BMA: t=0.6536, df=48.00, p=0.51651; IL-vHPC: t=3.940, df=48.00, **p=0.0010).

## Discussion

Both MeCP2 deficiency due to full knock-out of the gene, and MeCP2 duplication affect PV interneurons of transgenic mouse models [25–27], although this has never been addressed in the amygdala. Studies implicating the specific conditional knock-out of MeCP2 gene in PV interneurons further confirmed that the gene is crucial for their function in supporting plasticity and network activity patterns [8, 19]. Moreover, at the behavioral level, loss of the MeCP2 in PV interneurons replicates part of the symptoms related to Rett syndrome [7]. Mice with conditional Cre-dependent knock-out of MeCP2 in PV interneurons (PV-Cre::MeCP2^fl/y^) have altered social interaction behavior, fear learning, startle response and motor dysfunction [7]. It is crucial to understand which brain regions contribute to disease phenotypes related to alternations in MeCP2 gene. The motor dysfunction due to knock-out in the PV expressing neurons was found to be caused by cerebellum [28]. We suggested that the circuit abnormalities of PV-Cre::MeCP2^fl/y^, associated with the altered social interaction and fear learning, are likely to happen in the vHPC-BLA-mPFC triad with BLA as a hub and in BLA itself. Our data demonstrate that these mice have a deficit in the inhibitory input to the principal neurons in the lateral nucleus of the amygdala, as well as altered connectivity of BA with vHPC and mPFC. The data is important for future understanding of the PV-Cre::MeCP2^fl/y^ mice phenotype and the role of PV interneurons in the MECP2-related pathologies.

The microcircuit malfunction associated with general loss of MeCP2 in neurons has been demonstrated for mPFC and HPC [18, 27, 29, 30], but has never been addressed in BLA. The *MeCP2* knockdown targeted specifically at the BLA has been shown to impair amygdala-dependent learning and memory when compared with control mice [31]. The microcircuits in BLA, however, have never been studied in relation to MeCP2 loss. As the LA has a specific role in plasticity and is a key structure in fear conditioning, we concentrated on the local circuits involving PV interneurons in LA part of the amygdala. The reduction in the inhibition to the principal neurons in LA of PV-Cre::MeCP2^fl/y^ mice involved not only fast synaptic GABA_A_ receptor currents but also slow metabotropic inhibition by GABA_B_ receptor. GABA_B_ receptor activates GIRK channels in BLA, where GIRK1, GIRK2, and GIRK3 channel subunits mRNAs are shown to be expressed [32]. We used the high frequency stimulation in the LA to induce the synaptic release of GABA from the local interneurons. The release was not dependent on the LA innervation by cortical inputs, as the glutamatergic transmission was blocked with AMPA and NMDA antagonists. The reduction of both fast and slow inhibition was in line with observed decrease in the synaptic release of GABA from PV interneurons. There was no decrease in number of functional GABA_B_-receptors, as the potent agonist induced currents with comparable amplitudes in principal neurons of both genotypes. PV interneurons target perisomatic area of principal neurons in BLA and provide potent inhibition to regulate their excitability [33]. We observed elevated action potential firing of principal neurons in response to current steps in the LA of PV-Cre::MeCP2^fl/y^ mice, which is in line with the inhibitory reduction.

PV interneurons with MeCP2 loss in LA were receiving higher amplitudes of spontaneous excitatory currents but had unchanged inhibitory inputs. This suggests that MeCP2 affects glutamatergic transmission to PV interneurons, possibly by modulating postsynaptic AMPA-receptor functions, although we did not check it directly.

PV interneurons in BLA regulate the network oscillatory activities and network synchronization [34, 35]. The principal neurons of the BLA form reciprocal connections with vHPC and mPFC, where networks are also regulated by local PV interneurons. This connectivity is known to serve for social interaction behavior. As the PV-Cre::MeCP2^fl/y^ mice possess the knock-out in all PV-expressing neurons, it is impossible to claim one local population’s dominant role over another when resting state correlated neuronal activity is evaluated. In vivo recordings suggest that vHPC, BLA, and mPFC are wired through similar cellular patterns and that mPFC and vHPC recruit local interneurons in BLA. Within the BLA, the synchrony of LA and vHPC principal neurons firing is required for action potential generation in BA PNs [36]. We observed a significant reduction in BA functional connectivity with mPFC and vHPC in PV-Cre::MeCP2^fl/y^ mice. Moreover, we observed a reduction in correlated activity between mPFC and vHPC. In particular, the reduction was prominent between the vHPC and IL part of mPFC. Previous research has demonstrated that MeCP2 full knock-out or duplication disrupts oscillatory activity and synchronization in the mPFC [29, 37]. The vHPC projections to mPFC have been shown to malfunction in MeCP2 full knock-out mice [38]. In all these models, the circuit abnormalities were associated with impairment of social behavior. These data are in agreement with our findings and additionally support the idea of PV interneurons’ central role in social behavior divergence observed in animal models. Our data demonstrates that the functional connectivity is affected in the whole triad of these regions within the PV-Cre::MeCP2^fl/y^ mice’ brains in vivo. In addition to social behavior, this triad is involved in fear memory consolidation and extinction, where synchronized activity supported by reciprocal projections plays a significant role [39, 40]. Overall, the results point out that in the aim to better understand MeCP2 loss-related phenotype, the interplay between vHPC-BLA-mPFC should be taken into account.

## Materials and Methods

### Animals

B6.129P2-*Mecp2*^*tm1Bird*^/J (MeCP2^fl/fl^; Guy J, et al. 2001) mouse line was crossed with a B6.129P2-*Pvalb*^tm1(cre)Arbr/J^ (PV-Cre, JAX 008069) line to produce ablation of *Mecp2* gene selectively in the parvalbumin interneurons (PV-Cre::Mecp2^fl/y^) of males, their littermates with PV-Cre::Mecp2^x/y^ genotype further referred as PV-Cre were used as control. The mice were housed in individually ventilated cages with a 12-light/12-dark cycle (lights are off 7:00 p.m. – 7:00 a.m.), and food and water were supplied ad libitum. All animal experiments were performed following the University of Helsinki Animal Welfare Guidelines and approved by the National Animal Experiment Board of Finland (license numbers: KEK21-029, ESAVI/20432/2022). Adult animals were used for all experiments (1.5-2 months old).

### Viral injections

AAV viral vectors encoding for floxed EGFP (AAV8; pAAV-hSyn-DIO-EGFP; Addgene #50457) or floxed ChR2 (AAV1; pAAV-EF1a-double floxed-hChR2(H134R)-mCherry-WPRE-HGHpA; Addgene #20297) was injected bilaterally in the BLA of adult mice. Injections were done under anesthesia in a stereotaxic frame as described in Ryazantseva et al., 2020, using the following coordinates for BLA (from bregma): (1) AP −1,8 ML 3.4-3.5 DV 4.1–4.3 (2) AP −2.3 ML 3.4-3.5 DV 4.1–4.3.

### Electrophysiology

*Acute coronal sections* were prepared as previously (Englund et al., 2021). Briefly, the brain was removed and immediately placed in carbonated (95% O2/ 5% CO2) ice-cold N-Methyl-D-glucamine (NMDG) based protective cutting solution (pH 7.3–7.4) containing (in mM): 92 NMDG, 2.5 KCl, 1.25 NaH_2_PO_4_, 30 NaHCO_3_, 20 HEPES, 25 glucose, 2 thiourea, 5 Na-ascorbate, 3 Na-pyruvate, 0.5 CaCl_2_ and 10 MgSO_4_ (Ting et al., 2014). The vibratome (Leica VT 1200S) was used to obtain 300-µm-thick brain slices. Slices containing the amygdala were placed into a slice holder and incubated for 8–10 min in 34 °C in the NMDG–based solution. Slices were then transferred into a separate slice holder at room temperature with a solution containing (in mM): 92 NaCl, 2.5 KCl, 1.25 NaH_2_PO_4_, 30 NaHCO_3_, 20 HEPES, 25 glucose, 2 thiourea, 5 Na-ascorbate, 3 Na-pyruvate, 2 CaCl_2_ and 2 MgSO_4_ (saturated with 95% O_2_ /5% CO_2_). After 1–4 h of recovery, the slices were placed in a submerged heated (30-32 °C) recording chamber and continuously perfused with standard ACSF, containing (in mM): 124 NaCl, 3 KCl, 1.25 NaH_2_PO_4_, 26 NaHCO_3_, 15 glucose, 1 MgSO_4_ and 2 CaCl_2_.

*Whole-cell patch-clamp recordings* were done from amygdala neurons under visual guidance using glass capillary microelectrodes with resistance of 3–5.5 MΩ. Multiclamp 700B amplifier (Molecular Devices), Digidata 1322 (Molecular Devices) or NI USB-6341 A/D board (National Instruments), and WinLTP version 2.20 (Anderson et al., 2007) or pClamp 11.0 software were used for data collection, with low pass filter (10 kHz) and a sampling rate of 20 kHz. Whole-cell current clamp recordings of membrane excitability in principal neurons and PV interneurons in LA were performed using a filling solution containing (in mM): 135 K-gluconate, 10 HEPES, 5 EGTA, 2 KCl, 2 Ca(OH)_2_, 4 Mg-ATP, 0.5 Na-GTP, (280 mOsm, pH 7.2). The resting membrane potential was sampled and then adjusted to −70 mV. Depolarizing current steps with 500 ms (PV interneurons) or 1000 ms duration (principal neurons) were applied to induce action potential firing. The amplitude of the injected current was increased with 5-10 pA increments. Drug-induced GABA_B_-mediated and GIRK currents were recorded from LA PNs with the same solutions in the presence of antagonists for NMDA and AMPA receptors (50 µM AP-5 and 50 µM NBQX, respectively). 100 µM picrotoxin was included in the ACSF to block GABA_A_ receptors, and the holding potential of the neurons was −50 mV. A valve control system (VC-6-PINCH, Warner Instruments) with a manifold adjusted to the tube was used for fast drug application. ACSF, including the above antagonists, was applied directly to the LA to record a baseline before switching to a solution supplemented with the relevant drug (25 µM ML297, 25 µM SKF97541, or 2 mM Barium chloride). Stimulation-induced GABA_B_ postsynaptic responses (slow IPSC) recordings were done based on [21, 41], with a bipolar stimulating electrode placed at the apical cortical edge of the lateral amygdala (LA), while the principal neuron for the recording was approximately at the same depth in the middle of the lateral amygdala. Stimulation responses were evoked by a stimulus isolation unit (DS2A, Digitimer Ltd.) by applying a short burst of four stimuli at 100 Hz in the presence of picrotoxin, NBQX, and AP-5 at the holding potential of −50 mV. The stimulation power was 28-32 V and was adjusted in this range to reach maximum response amplitudes. After the stimulation was not able to induces larger current, the V value was used for stimulation and recording of series of responses. The GABA_B_ antagonist 5 μM CGP 55845 was added to the recording chamber through the main ASCF flow. Spontaneous synaptic currents (sEPSCs and sIPSCs) in PV interneurons and LA principal neurons were recorded under a whole-cell voltage-clamp with a filling solution containing (in mM): 135 K-gluconate, 10 HEPES, 5 EGTA, 2 KCl, 2 Ca(OH)_2_, 4 Mg-ATP, 0.5 Na-GTP, at holding potential of −50 mV. For light-induced IPSCs, recordings were done using a high-chloride intracellular solution containing (in mM): 130 CsCl, 10 HEPES, 0.5 EGTA, 8 NaCl, 4 Mg-ATP, 0.3 Na-GTP, 5 QX314 (280 mOsm, pH 7.2), at −70 mV holding potential in the presence of NBQX and AP-5. Optical stimulation of PV interneurons was done using pE-2 LED system (CoolLED). Short pulses (0.5 ms, 50 ms interval) of 470 nm blue light were used for paired-pulse stimulation. WinLTP software (Anderson and Collingridge, 2007) was used to calculate the peak amplitude of the evoked synaptic responses. For analysis of paired-pulse ratio (PPR), 7–10 responses were averaged in each experimental condition. PPR was calculated as the amplitude ratio of response 2/response 1. The frequency and amplitude of spontaneous synaptic events were analyzed using miniAnalysis program 6.0.3. sIPSCs and sEPSCs were identified in the analysis as outward or inward currents (for IPSCs depending on experimental conditions) with typical kinetics, respectively, that were at least 3 times the amplitude of the baseline level of noise. For the pooled data, averages for baseline were calculated over a 10-minute period. Action potential frequencies were analyzed using the threshold search algorithm in Clampfit software. AP half-width potential was analyzed from the 3rd spike in the train using Clampfit software.

### Confocal microscopy and spine density analysis

Biocytin (HelloBio) was added to the recording solution. For post hoc morphological characterization of biocytin-filled neurons, slices were fixed overnight in a 4% paraformaldehyde (4°C), after which they were washed with cold phosphate-buffered saline (PBS) and permeabilized with 0.3% Triton-X 100 (Sigma-Aldrich) in PBS for overnight at 4°C. Alexa Fluor 568 streptavidin (1:500; Life Technologies) was added to the permeabilization solution and incubated for overnight at 4°C. PBS-washed slices were mounted onto slides and blind-coded for morphological analysis. Dendritic spines were imaged using an LSM Zeiss 710 confocal microscope (Zeiss alpha Plan146 Apochromat 63x/1.46 OilKorr M27 objectives). Spines were imaged with a resolution of 15.17 pixels/μm and a Z-stack interval of 0.5 μm. Spine density was evaluated on the secondary dendrites. Actual spine detection was done using the NeuronStudio software to quantify spines in a Z-stack image. Verification of the spine detection was done semi-manually. Values of spine density per cell were used for statistical analysis.

### In vivo Functional ultrasound

Resting-state functional connectivity between the medial prefrontal cortex (mPFC), basolateral amygdala (BLA) and ventral hippocampus (vHPC) in littermates of 3 litters using Iconeus One functional ultrasound imaging system. The animals were anesthetized using medetomidine (1 mg/kg) + ketamine (75 mg/kg), after which the head fur was shaved. The animal head was fixed in a stereotaxic frame with heat pad (35°C) and ultrasound gel was spread on the scalp. The probe was controlled using the IcoScan (v. 1.3.1) software, first placing it near the scalp and then locating the posterior cerebral arteries (PCA). A coronal mapping (Angio3D) scan was performed from the PCA to the mPFC with a slice interval of 0.1 mm. The scan was uploaded to IcoStudio (v. 1.2.1) software and was mapped using the automated mapping system in the software. Regions of interest were selected (mPFC, BLA, vHPC) and markers were placed on the mapping. Probe coordinates were computed based on these markers, to align all areas of interest in a single plane. The coordinates were manually uploaded to the IcoScan software, and a 20-minute 2D functional ultrasound scan was performed. After the scan, the animals were placed into a heat chamber (35°C) and injected with atipamezole (0.5 mg/kg) for recovery. IcoStudio software was used to compute a functional connectivity matrix, based on the functional ultrasound scan. Baseline correction and a 0.2Hz low pass filter were applied to the final results. For analysis, the matrix values (Pearson’s correlation coefficients) were Fisher transformed.

## Acknowledgements

Authors thank Janne Sulku, the personnel in the Laboratory Animal Center and In Vivo Animal Imaging Platform (HAIP) of the University of Helsinki for expert technical help. This study was financially supported by the Academy of Finland (#330710, S.E.L.; #330298, M.R., #350193, T.A.), Sigrid Juselius Foundation (#1158, S.E.L.), Jane and Aatos Erkko Foundation (S.M., Finland). In Vivo Animal Imaging Platform (HAIP) is supported by Euro-BioImaging Research infostructure.

## Notes

### Competing Interest Statement

The authors have declared no competing interest.

### Summary of Updates

Unnecessary data that were out of the paper scope was removed, text was improved

